# Expression of *TaTAR2.3-1B*, *TaYUC9-1* and *TaYUC10* correlates with auxin and starch content of developing wheat grains

**DOI:** 10.1101/2020.10.12.336560

**Authors:** Muhammed Rezwan Kabir, Heather M. Nonhebel, David Backhouse, Gal Winter

**Author notes:** Corresponding author: Heather. M. Nonhebel.

## Abstract

The role of auxin in developing grains of wheat (*Triticum aestivum*) is contentious with contradictory reports indicating either positive or negative effects of IAA (indole-3-acetic acid) on grain size. In addition, the contributions to the IAA pool from de novo synthesis via tryptophan, and from hydrolysis of IAA-glucose are unclear. Here we describe the first comprehensive study of tryptophan aminotransferase and indole-3-pyruvate mono-oxygenase expression during wheat grain development from 5 to 20 days after anthesis. A comparison of expression data with measurements of endogenous IAA via combined liquid chromatography-tandem mass spectrometry with heavy isotope labelled internal standards indicates that TaTAR2.3-1B, TaYUC9-A1, TaYUC9-B, TaYUC9-D1, TaYUC10-A and TaYUC10-D are primarily responsible for IAA production in developing grains. Furthermore, we show that IAA synthesis is controlled by genes expressed specifically in developing wheat grains as has already been reported in rice (*Oryza sativa*) and maize (*Zea mays*). Our results cast doubt on the proposed role of *THOUSAND-GRAIN WEIGHT* gene, *TaTGW6*, in promoting larger grain size via negative effects on grain IAA content. The work on *TaTGW6* has overlooked the contribution of the dominant IAA biosynthesis pathway. Although IAA synthesis occurs primarily in the endosperm of wheat grains, we show that the *TaYUC9-1* group is also strongly expressed in the embryo. Within the endosperm, *TaYUC9-1* expression is highest in aleurone and transfer cells, supporting data from other cereals suggesting that IAA has a key role in differentiation of these tissues.

## Introduction

Global wheat (*Triticum aestivum*) production has reached 760.1 million tonnes (FAOSTAT 2020); however, a substantial increase in yield from the existing land area is essential to meet the needs of the rapidly increasing world population. The International Wheat Yield Consortium (WYC) has introduced a strategy to improve yield potential that could accelerate breeding of high yielding wheat varieties (Foulkes et al. 2011). This has identified grain number and grain weight as two key yield-determining factors. The plant hormone auxin or IAA (indole-3-acetic acid) appears to play a major role in both of these aspects of grain development.

In particular, inactive alleles of two genes *TaTGW6* and *TaTGW-7A* are reported to improve grain size in wheat via negative effects on the grain IAA content (Hu et al. 2016a; Hu et al. 2016b). *TaTGW6* is homologous to the rice gene *TGW6,* reported by Ishimaru et al. (2013) to encode an IAA-glucose hydrolase. Although the work on wheat did not characterise the gene product of *TaTGW6,* Hu et al. (2016a) reported that an inactive allele, *TaTGW6-c,* as well as a mutant allele, *TaTGW6-b,* were associated with lower IAA content of grains at 20 and 30 days after anthesis (DAA) as well as higher grain weight. *TaTGW-7A* is reported as an indole-3-glycerol phosphate synthase (IGPS) like gene (Hu et al. 2016b). Again, there was no characterisation of the gene product. However, the authors reported a similar lower IAA content in grains with the inactive *TaTGW-7Aa* allele as that found in *TaTGW6-b* and *TaTGW6-c* plants.

In contrast, the majority of relevant publications report a positive involvement of IAA on grain size in a number of cereals. For example, the shrunken grain phenotype of defective endosperm and defective kernel mutants of maize, *de18* and *dek18* appears to result from low levels of IAA in the developing grains caused by mutations affecting *ZmYUC1* expression (Bernardi et al. 2012; Bernardi et al. 2016). In rice, a mutation in the IAA biosynthesis gene *OsTAR2/FIB* is associated with the *tillering and small grain 1 (tsg1)* phenotype (Guo et al. 2019). Finally, in wheat, Shao et al. (2017) showed that overexpression of *TaTAR2.1-3A* increases IAA accumulation in grains and enhances plant height, spike number, biomass and grain yield. Thus the claim that *TaTGW6* and *TaTGW-7A* have a positive effect on grain size via a reduction in IAA content requires careful scrutiny.

In plants, including developing seeds, IAA is produced primarily from tryptophan via the actions of tryptophan aminotransferase of Arabidopsis1 (TAA1) and its related proteins (TAR), and indole-3-pyruvate monooxygenase known as YUCCA (Mashiguchi et al. 2011; Won et al. 2011). In assuming that inactivation of a glucose hydrolase gene can lead to a large reduction in grain IAA content, the *TGW6* work has overlooked this major source of IAA. On the other hand, the same authors suggest that *TaTGW-7A* affects grain IAA content via its effects on tryptophan production. The TAR/YUCCA pathway is known to occur in wheat grains from three studies. Shao et al. (2017) reported a comprehensive study of the *TAR* gene family based on an early version of the wheat genome and showed that of all *TAR* genes, *TaTAR2.3* was maximally expressed in grains. However, they did not investigate the timing of *TaTAR2.3* expression during grain fill or present any data on the IAA content of grains. Li et al. (2014) showed *TaYUC10.3* is highly expressed in developing wheat grains, with expression increasing throughout the grain-fill period but did not investigate any other YUCCA genes from wheat or relate gene expression to the IAA content. Finally, Tuan et al. (2019) described expression changes in “*TaTAR2”* (actually *TaTAR2.3-1D*) and “*TaYUC11*” from microarray data during seed maturation (20-50 DAA) and also measured the IAA content of endosperm and embryo tissue. However, there have been no comprehensive studies of the *YUCCA* gene family in wheat, their expression profile during grain development or correlation with the grain IAA content.

The aim of our work was therefore to evaluate the role of the TAR/YUCCA pathway in the regulation of IAA content during grain fill in wheat. The study comprised the first compressive phylogenetic and expression analysis of all *TAR* and *YUCCA* genes in wheat. The expression profile of major IAA biosynthesis genes during early grain development was compared with changes in the grain IAA content measured by the most accurate and specific method available, as well as with changes in the starch content of grains. To inform future studies on the specific signalling role(s) of IAA in wheat grains, we also investigated the location of IAA production via dissection of grains into embryo and endosperm fractions as well as by mining information from published RNA-sequencing (RNA-seq) studies.

## Materials and methods

### Data mining and Bioinformatic analysis

Protein sequences of all *TAR* and *YUCCA* genes from wheat (*Triticum aestivum*) (IWGSC RefSeq v1.1), rice (*Oryza sativa*), brachypodium (*Brachypodium distachyon*) and barley (*Hordeum vulgare*) were downloaded from EnsemblPlants 47 (Kersey et al. 2015) following BlastP searches using the query sequences OsTAR2, OsYUC1 and OsYUC9 from rice. Arabidopsis TAA1 and TAR sequences were downloaded from The Arabidopsis Information Resource (TAIR) (Rhee et al. 2003). Phylogenetic analyses of the protein sequences were carried out in MEGA7.0.26 (Kumar et al. 2016) using the Maximum Likelihood method (Jones et al. 1992). Multiple sequence alignments (MSA) were performed by MUSCLE (Edgar 2004). Bootstrap confidence levels were obtained using 500 replicates (Felsenstein 1985). Evolutionary distances were computed using Poisson correction method (Zuckerkandl and Pauling 1965). RNA-seq data available in expVIP (http://www.wheat-expression.com) (Borrill et al. 2016; Ramírez-González et al. 2018) were used for global evaluation of gene expression, as well as for investigating gene expression in dissected aleurone and transfer cell tissues.

### Plant materials

Wheat plants (Chinese Spring) were grown in 20 cm × 20 cm pots under natural light at 23/14°C (day/night temperatures) in the glasshouse at the University of New England, Armidale, NSW. Pots were fertilized once a week with Thrive^®^ (Yates, 1 g/L) from the initiation of tillering stage. Spikes were tagged when the first spikelet reached anthesis. Grain samples (70–90 mg) were harvested at the same time (4:00 to 5:00 pm) each day 5, 10, 15 and 20 DAA, then snap-frozen in liquid nitrogen and stored at −80°C. Some 15 DAA grains were manually dissected into embryos and endosperms. Grain samples from individual spikes were kept separate and all analyses were done on at least three independent biological replicates harvested from different plants, on different days.

### RNA extraction and quantitative analysis

Wheat grain samples, harvested and stored as described above, were ground in liquid nitrogen and total RNA was extracted using Trizol (Invitrogen). RNA concentration and 260/280 ratio were determined using a NanoDrop ND-8000 Spectrophotometer (Thermo Scientific). RNA samples with A_260/280_ of 1.8–2.0 were used for further analysis. The RNA quality was checked by agarose gel electrophoresis for two clear bands of 18S and 28S rRNAs (Nolan et al. 2006). Primers specific for each group of three paralogues on the A, B and D genomes were designed with melting points in the range of 58–60°C and product sizes between 108–251 bp (Supplementary Table S1). Initial amplification was carried out using a One-Step RT-PCR Kit (QIAGEN) in a BIO-RAD T100 Thermal Cycler, with gel analysis to confirm a single product of the expected size. cDNAs were prepared using the SensiFAST cDNA Synthesis Kit (Bioline). Quantitative real-time PCR reactions using cDNA samples as template were carried out using a SensiFAST SYBR No-ROX Kit (Bioline) in a CFX96 Touch (BIO-RAD) machine following manufacturer’s instructions. Negative control reactions without reverse transcriptase as well as reactions with no template were included. Three biological and three technical replicates were carried out for each primer set. The qRT-PCR program included 40 cycles of 95°C 3 min, 95°C 10 s, 55°C 30 s and 72°C 5 s. Expression was calculated relative to two reference genes, translation elongation factor EF-I alpha Ta53964 (Paolacci et al. 2009) and a cyclin-like protein Ta27922 (Wu et al. 2015). Expression data presented are the average and standard error of biological replicates. Amplified products were sent to the Australian Genome Research Facility for sequencing to confirm identity following the use of the Wizard SV Gel and PCR Clean-Up System (Promega).

### IAA extraction and analysis

Wheat grain samples (70–90 mg) were ground in liquid nitrogen; 200 μL of 65% isopropanol /35% 0.2 M pH 7.0 imidazole buffer (Chen et al. 1988) was added with [^13^C_6_] IAA internal standard (Cambridge Isotope Laboratories Inc.), and samples were extracted on ice for 1 h. Amounts of standards added varied with the age of samples to ensure that the concentration of standards was similar to that of endogenous IAA; 78 ng of [^13^C_6_] IAA was added to 10, 15 and 20 DAA samples; 16 ng of [^13^C_6_] IAA was added to 5 DAA samples. Blank samples without plant tissue were taken through the entire extraction and analysis protocol to ensure that no contamination from the unlabelled IAA in the laboratory occurred.

Following extraction, samples were diluted with 2 mL deionized water, centrifuged and the supernatant transferred to a glass tube. Sample clean-up followed the solid-phase extraction (SPE) protocol of (Barkawi et al. 2008) with minor modifications. Samples prepared as above were added to discovery^®^ DSC-NH2 SPE 50 mg/mL tubes (Supelco) that had been prewashed sequentially with 500 μL hexane, 500 μL acetonitrile, 500 μL water, 500 μL 0.2 M pH 7.0 imidazole buffer and 4.5 mL deionized water. After loading each sample, SPE columns were washed sequentially with 500 μL each of water, hexane, ethyl acetate, acetonitrile, methanol and 600 μL 0.25% phosphoric acid. The IAA was eluted in 1.8 mL 0.25% phosphoric acid and the pH of the eluate was adjusted to 3.0–3.5 with 150 μL 0.1 M, pH 6.0 succinate buffer. This fraction was added to Strata-X SPE (8B-S100-UBJ 60 mg/3 mL; Phenomenex) that had been prewashed with 1 mL hexane, 1 mL methanol and 2 mL water. Samples loaded on the SPE columns were washed with 3×1 mL water and 100 μL acetonitrile before eluting the IAA in 1 mL acetonitrile.

Samples from SPE clean-up were stored at −20°C. Immediately prior to analysis, they were reduced to dryness under a stream of N_2_ and redissolved in 20 μL acetonitrile and 80 μL 0.01 M aqueous acetic acid. The analysis of ^12^C:^13^C IAA was conducted using a triple quadrupole Liquid Chromatograph Mass Spectrometer (LCMS)-8050, (Shimadzu) with XBridge™ C18 3.5 μm, 2.1×50 mm column (Phenomenex). The chromatography solvent was 20% acetonitrile: 80% 0.01 M acetic acid at a flow rate of 0.2 mL/min. The nebulizing, heating and drying gas flow were 3 L/min, 10 L/min and 10 L/min, respectively. Interface temperature was 300°C, DL was 250°C and the heat block temperature was 400°C. The interface used a capillary voltage of 4 kV. The mass spectrometer was operated in multiple-reaction-monitoring mode (collision energy, 14.0 eV), transitions from *m/z* 174.10 to 130.10 for [^12^C_6_] and *m/z* 180.20 to 136.15 for [^13^C_6_] were monitored. A series of standard mixtures of [^13^C_6_] and unlabelled IAA in different ratios 10:1 to 1:10 were also assayed to confirm accuracy of quantitative analysis. Data were obtained from the average of two technical replicates first, then the average and the standard error of three biological replicates from each developmental stage.

### Starch assay

Frozen wheat grain samples were freeze-dried then placed in microfuge tubes with stainless steel beads and ground using a TissueLyser II (QIAGEN) for 3 min at a frequency of 30/s. Starch extraction and analysis followed the methods of Zhao et al. (2010). After the extraction of soluble sugars in 80% ethanol, the remaining starch pellet was hydrolysed sequentially with amylase (Sigma A3403) and amyloglucosidase (Sigma A7095). The resulting glucose was assayed using glucose HK assay reagent (Sigma G3293) in microtitre plates using a SPECTROstar^Nano^ (BMG LABTECH) at 340 nm. Three biological replicates were extracted for each time period; three technical replicates were assayed for each sample.

## Results

### Wheat has orthologues of OsTAR1 and OsTAR2 as well as a wheat-specific branch of TaTAR2

A BlastP search of the wheat proteome found 15 co-orthologues of Arabidopsis TAR2. Their encoding genes are listed in Table 1, with IWGSC RefSeq v1.1 gene IDs. Twelve of these were previously named by Shao et al. (2017) as *TaTAR2.1* to *TaTAR2.5* with suffixes indicating chromosome/genome. We followed a similar format, naming the three additional genes as group *TaTAR2.6*. Two of these are tandem repeats designated as *TaTAR2.6-1Ba* and *TaTAR2.6-1Bb* whereas the chromosomal location of the third *TaTAR2.6-U* is currently unknown. *TaTAR2.1, TaTAR2.2* and *TaTAR2.3* have one copy in each of the A, B and D genomes. On the other hand, *TaTAR2.4*, *TaTAR2.5* and *TaTAR2.6* do not have a copy on the D genome.

**Table 1.**
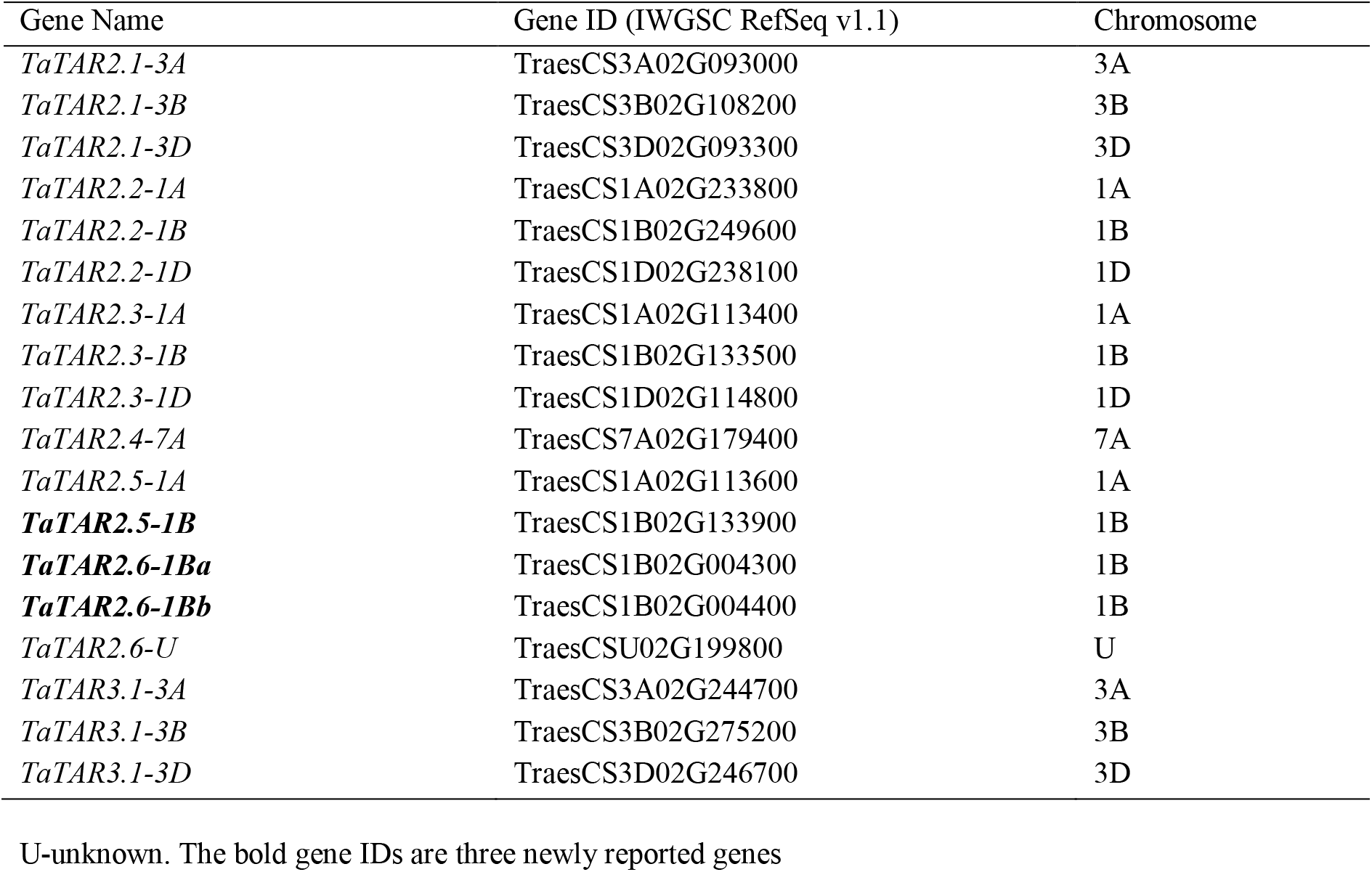
List of *TaTAR* genes in the wheat genome

The protein phylogenetic tree in Fig. 1 compares wheat TAR sequences with rice and Arabidopsis enzymes that have demonstrated tryptophan aminotransferase activity. The tree also has a branch containing three TaTAR3 proteins orthologous to OsTAR3. These are usually designated as alliinases, and unlikely to be involved in IAA production. Three proteins designated as TaTAR2.3-1A/B/D are co-orthologues of OsTAR1/OsFBL in rice. Barley and Brachypodium also have at least one protein in this clade. A second clade with a high bootstrap value contains OsTAR2/OsFIB as well as six closely related wheat proteins, TaTAR2.1-3A/B/D and TaTAR2.2-1A/B/D and one protein each from barley and Brachypodium. The remaining six wheat proteins have been placed in the same clade as OsTAR1 and TaTAR2.3 but with a low bootstrap value indicating ambiguity. This branch has no proteins from the other cereals.

**Fig. 1.**
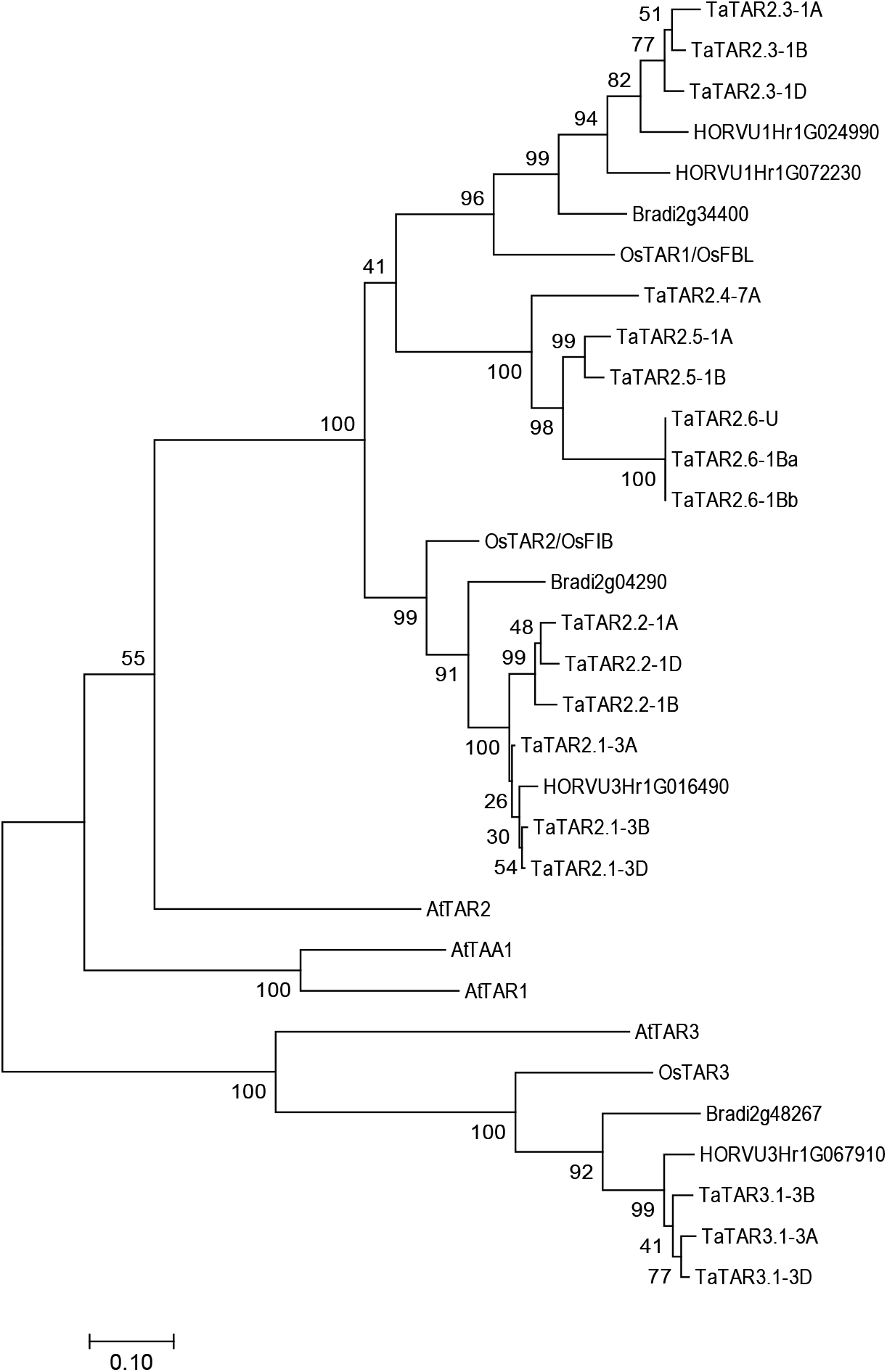
Phylogenetic tree showing relationships between TAR proteins from *Triticum aestivum* (TaTAR), *Oryza sativa* (OsTAR), *Brachypodium distachyon* (Brad), *Hordeum vulgare* (HORVU) and *Arabidopsis thaliana* (At). The tree was constructed in MEGA7.0.26 (Kumar et al., 2016) using Maximum Likelihood method (Jones et al., 1992). Multiple sequence alignment was performed by MUSCLE (Edgar, 2004). Bootstrap confidence levels were obtained using 500 replicates (Felsenstein, 1985). Evolutionary distances were computed using Poisson correction method (Zuckerkandl and Pauling, 1965). Scale bar=0.10 amino acid substitutions per site

### Expression of *TaTAR2.3-1B* in grains increases from 5 to 15 DAA

We investigated the expression of all *TaTAR2* genes at four times during wheat grain fill, 5, 10, 15 and 20 DAA. Due to the large number and high similarity of genes from each genome, six primer sets were designed, each to amplify all genes from sets *TaTAR2.1* to *TaTAR2.6* as shown in Supplementary Table S1. Initial RT-PCR analysis showed efficient amplification of a single band of the expected size for the *TaTAR2.3* group only. In contrast, *TaTAR2.1*, *TaTAR2.2* and *TaTAR2.5* groups showed very low amplification and there was no amplification of *TaTAR2.4* and *TaTAR2.6* groups. We therefore investigated the expression of *TaTAR2.3* by qRT-PCR. A large (44-fold) up-regulation occurred between 5 and 15 DAA after which expression appeared to plateau (Fig. 2a). The location of *TaTAR2.3* expression within grains at 15 DAA was also investigated by dissection of the grains into embryo and endosperm components (Fig. 2b). This revealed endosperm-specific gene activity. Sequencing of the PCR product from 15 DAA samples indicated the B genome copy of *TaTAR2.3* was the primary gene expressed.

**Fig. 2.**
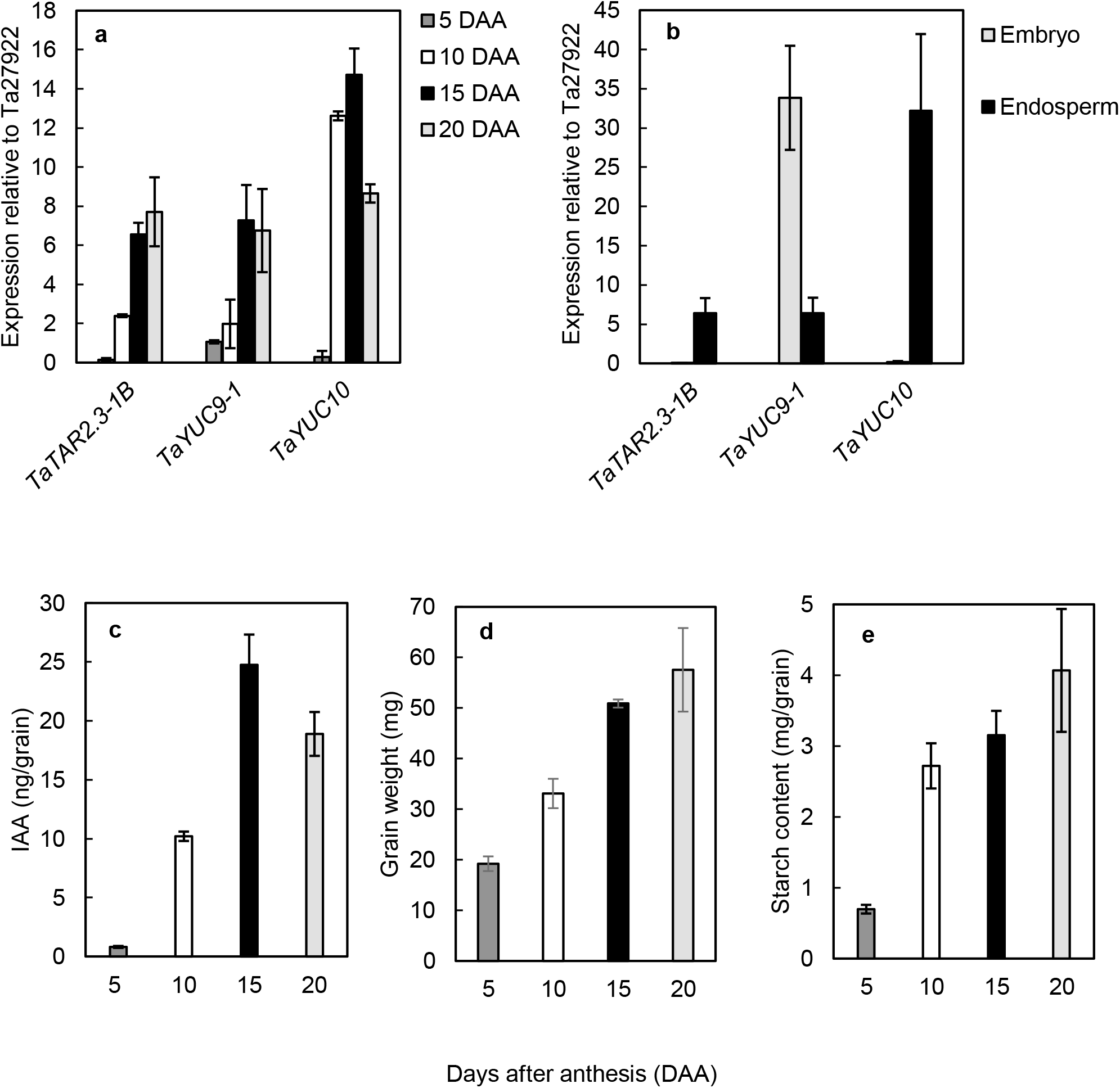
Quantitative RT-PCR analysis of relative transcript abundance for TaTAR2.3, TaYUC9-1 and TaYUC10 during grain development, whole grains (a), in dissected embryos and endosperms at 15 DAA (b), IAA content in ng per grain measured by LC-MS/MS MRM with [13C] IAA as the internal standard (c), grain fresh weight (d) and starch content (e). Expression relative to reference gene Ta27922 is shown, expression relative to the second reference gene Ta53964 showed the same trend. All data points represent the mean of three biological replicates ± the standard error of the means

Additional information on *TaTAR2* expression, from RNA-seq data in the expVIP database is shown in Fig. 3. This confirmed *TaTAR2.3-1B* is the most highly expressed gene in wheat grains, with maximum up-regulation between 10 and 20 DAA. The D genome copy of *TaTAR2.3* was also expressed but at much lower levels. Results from manually dissected tissue layers (Pfeifer et al. 2014), indicated that expression occurred across all parts of the endosperm including starchy endosperm, aleurone layer and transfer cells. In addition, *TaTAR2.5-1B*, *TaTAR2.2-1A* and *TaTAR2.2-1D* are expressed very early in grain development at 2 DAA, followed by a decrease in expression.

**Fig. 3.**
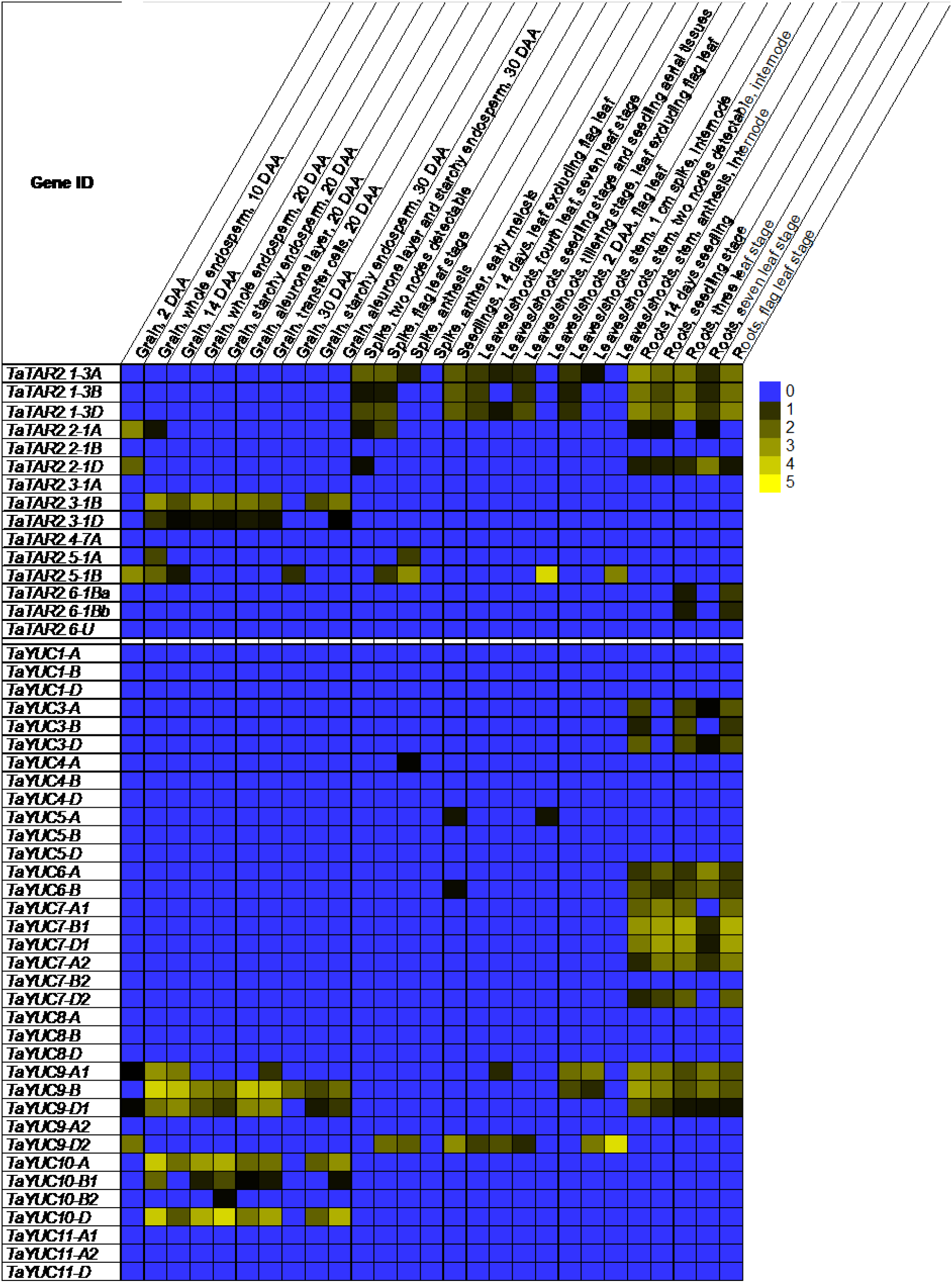
Heat map depicting expression of *TaTAR2* and *TaYUC* genes from RNA-seq data in expVIP. Data were taken from different studies using the Chinese Spring variety. The relative expression values are normalized as tpm (transcripts per million)

### Comprehensive phylogenetic analysis of YUCCA proteins and systematic naming of wheat YUCCAs

The wheat genome contains 35 members of the YUCCA gene family. These are listed in Table 2 along with their chromosomal location and IWGSC RefSeq v1.1 gene IDs. As there has been no previous comprehensive study of the YUCCA gene family in wheat, we have named these systematically according to their homology with rice *YUCCA* genes following naming structure in the Wheat Gene Catalogue (https://wheat.pw.usda.gov/GG3/wgc). We suggest renaming the TaYUC10 group, previously designated as *TaYUC10.1, TaYUC10.2* and *TaYUC10.3* by Li et al. (2014) as *TaYUC10-B1, TaYUC10-A* and *TaYUC10-D*, respectively. In addition, Tuan et al. (2019) referred to TraesCS5B02G216000 as *TaYUC11* by homology to Arabidopsis *YUC11*. We suggest it is more logical to refer to this gene as *TaYUC9-B* as it is orthologous to *OsYUC9,* and there are other wheat genes orthologous to *OsYUC11*.

**Table 2.** List of *TaYUC* genes in the wheat genome

| Gene Name | Gene ID (IWGSC RefSeq v1.1) | Chromosome |
| --- | --- | --- |
| *TaYUC1-A* | TraesCS3A02G232600 | 3A |
| *TaYUC1-B* | TraesCS3B02G261900 | 3B |
| *TaYUC1-D* | TraesCS3D02G220500 | 3D |
| *TaYUC3-A* | TraesCS3A02G280900 | 3A |
| *TaYUC3-B* | TraesCS3B02G314700 | 3B |
| *TaYUC3-D* | TraesCS3D02G280900 | 3D |
| *TaYUC4-A* | TraesCS3A02G149500 | 3A |
| *TaYUC4-B* | TraesCS3B02G176800 | 3B |
| *TaYUC4-D* | TraesCS3D02G157600 | 3D |
| *TaYUC5-A* | TraesCS5A02G102700 | 5A |
| *TaYUC5-B* | TraesCS5B02G107000 | 5B |
| *TaYUC5-D* | TraesCS5D02G114500 | 5D |
| *TaYUC6-A* | TraesCS4A02G313200 | 4A |
| *TaYUC6-B* | TraesCS5B02G566700 | 5B |
| *TaYUC7-A1* | TraesCS2A02G011500 | 2A |
| *TaYUC7-B1* | TraesCS2B02G010100 | 2B |
| *TaYUC7-D1* | TraesCS2D02G012100 | 2D |
| *TaYUC7-A2* | TraesCS2A02G533200 | 2A |
| *TaYUC7-B2* | TraesCS2B02G562800 | 2B |
| *TaYUC7-D2* | TraesCS2D02G535100 | 2D |
| *TaYUC8-A* | TraesCS4A02G027500 | 4A |
| *TaYUC8-B* | TraesCS4B02G278300 | 4B |
| *TaYUC8-D* | TraesCS4D02G276600 | 4D |
| *TaYUC9-A1* | TraesCS5A02G217200 | 5A |
| *TaYUC9-B* | TraesCS5B02G216000 | 5B |
| *TaYUC9-D1* | TraesCS5D02G225200 | 5D |
| *TaYUC9-A2* | TraesCS7A02G065600 | 7A |
| *TaYUC9-D2* | TraesCS7D02G060000 | 7D |
| ***TaYUC10-A*** | TraesCS4A02G325400 | 4A |
| ***TaYUC10-B1*** | TraesCS5B02G538300 | 5B |
| *TaYUC10-B2* | TraesCS6B02G033700 | 6B |
| ***TaYUC10-D*** | TraesCS5D02G556600 | 5D |
| *TaYUC11-A1* | TraesCS4A02G373500 | 4A |
| *TaYUC11-A2* | TraesCS7A02G075400 | 7A |
| *TaYUC11-D* | TraesCS7D02G070900 | 7D |

Genes have been numbered by homology to rice genes. Genes in bold were previously reported as *TaYUC10.2*, *TaYUC10.1* and *TaYUC10.3*, respectively by Li et al. (2014).

Figure 4 shows the phylogenetic relationships between wheat and rice YUCCA proteins. The tree comprises two major clades; the larger clade I contains OsYUC1-8 as well as orthologous wheat proteins. OsYUC2 is missing from the tree as *OsYUC2* appears not to be expressed in rice and it also has no wheat orthologues. Most of the TaYUC sequences in clade I are encoded by each of the A, B and D genomes. However, TaYUC-6 is found in the A and B genomes only. The smaller clade II contains OsYUC9, OsYUC11, OsYUC12 and OsYUC14 (*OsYUC10* and *OsYUC13* being probable pseudogenes) as well as 12 wheat proteins. Wheat has five proteins in the OsYUC9 branch. Three of these, TaYUC9-A1, TaYUC9-B and TaYUC9-D1 are co-orthologous with OsYUC9 and we refer to these are the TaYUC9-1 group. Two less similar proteins in this branch have been named as TaYUC9-A2 and TaYUC9-D2. Wheat also has three co-orthologues of OsYUC11; TaYUC11-A1, TaYUC11-D and TaYUC11-A2. In addition, there is a group of four proteins designated as TaYUC10, three of which were previously reported by Li et al. (2014). TaYUC10 proteins have highest amino acid similarity to OsYUC11 but are also similar to OsYUC12/14; this agrees with their positioning in the tree. Due to their importance in grain development, the relationship between proteins in clade II was further investigated with the addition of homologous proteins from Brachypodium and barley (Supplementary Figure S1). This revealed that both species have at least one orthologue of OsYUC9 and OsYUC11 but the OsYUC12/14 branch does not have orthologues in wheat, barley or Brachypodium, all members of the Pooideae. Instead, these cereals have a separate branch including TaYUC10, which has undergone considerable gene duplication within each species, particularly barley.

**Fig. 4.**
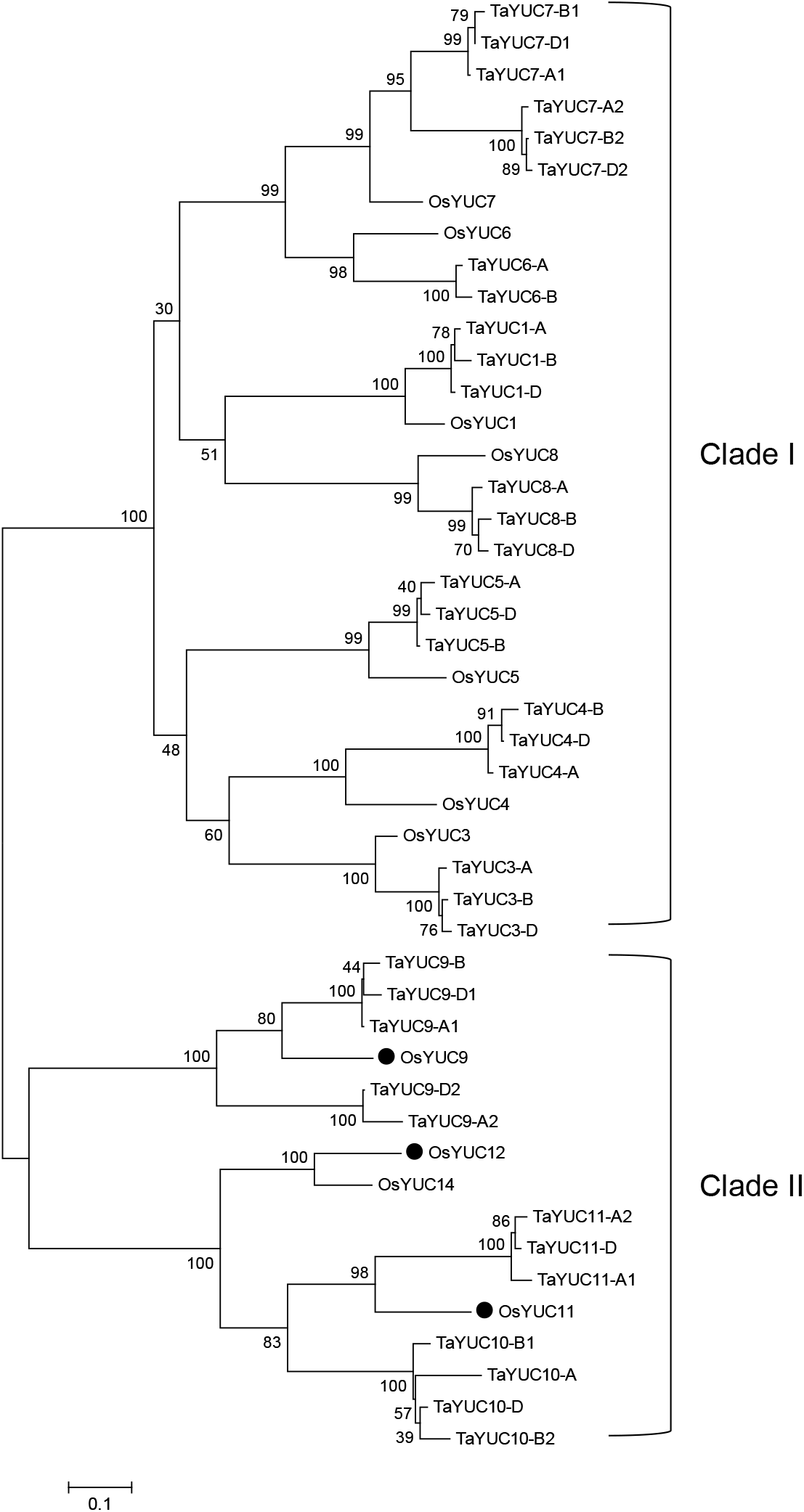
Phylogenetic tree showing relationships between YUCCA proteins from *Triticum aestivum* (TaYUC) and *Oryza sativa* (OsYUC). The tree was constructed in MEGA7.0.26 (Kumar et al. 2016) using Maximum Likelihood method (Jones et al. 1992). Multiple sequence alignment was performed by MUSCLE (Edgar 2004). Bootstrap confidence levels were obtained using 500 replicates (Felsenstein 1985). Evolutionary distances were computed using Poisson correction method (Zuckerkandl and Pauling 1965). Scale bar=0.10, amino acid substitutions per site. Black dots are encoded by genes expressed in rice grains

### Expression of *TaYUC9-1* and *TaYUC10* in developing grains increases from 5 to 15 DAA

Figure 3 summarises information from RNA-seq data (expVIP database), on the expression throughout the plant as well as during grain development of *TaYUC* genes. None of the *TaYUC* genes in clade I has significant expression in wheat grains. Some expression occurs in leaves and the spike at anthesis. However, the greatest expression of genes in this clade is in the roots, where *TaYUC3, TaYUC6* and *TaYUC7* are the most highly expressed gene groups, with expression of genes from all three genomes. Conversely, a number of clade II genes show very high expression in grains. Greatest up-regulation is found between 10 and 20 DAA, with *TaYUC9-A1*, *TaYUC9-B*, *TaYUC9-D1*, *TaYUC10-A* and *TaYUC10-D* being the genes with highest activity. A different gene, *TaYUC9-D2* has high expression at 2 DAA in grains, after which it is down-regulated, with no observable expression at 10 DAA. Expression of the *TaYUC10* group is restricted to the grains. However, the *TaYUC9-1* group is also active in vegetative tissue, particularly the stem and roots. *TaYUC9-D2* is the gene with highest activity in leaves and spike.

As gene expression in the developing grains was the primary focus of this study, we carried out a quantitative expression analysis of *TaYUC* genes in clade II from 5 to 20 DAA. Primers were designed to amplify genes from all three genomes as shown in Supplementary Table S1. Initial RT-PCR screening demonstrated successful amplification of the *TaYUC9-1* and *TaYUC10* groups, producing a single product of the correct size. On the other hand, no product was obtained for *TaYUC9-2* and *TaYUC11* confirming that these genes have very little or no expression in grains from 5 to 20 DAA. Investigation by qRT-PCR showed strong up-regulation of the *TaYUC9-1* and *TaYUC10* groups during grain development, as shown in Fig. 2a. The *TaYUC10* genes were up-regulated earlier than the *TaYUC9-1* group, reaching maximum expression at 10 DAA and decreasing activity at 20 DAA. In contrast, the expression of *TaYUC9-1* was primarily up-regulated between 10 and 15 DAA, and showed no reduction in expression at 20 DAA. The location of expression of *TaYUC9-1* and *TaYUC10* groups was also investigated following dissection of wheat grains into the embryos and endosperms. The results in Fig. 2b suggest *TaYUC9-1* genes are expressed in both tissues with higher expression in embryos, whereas *TaYUC10* genes are only expressed in endosperm. Examination of the RNA-seq data from dissected tissue layers provides further information on localisation of *TaYUC* gene expression. These data indicate that highest expression of *TaYUC9-1* occurs in the aleurone and transfer cells, whereas *TaYUC10* has the highest expression in the starchy endosperm. Nevertheless, expression of both genes appears to occur throughout the endosperm. No RNA-seq data is available on expression in wheat embryos of the Chinese Spring variety. However, data from the variety Azhurnaya (not shown) confirm our observations of expression of *TaYUC9-1* but not *TaYUC10* in this tissue.

Sequencing of the RT-PCR products was used to clarify which copy(s) of the *TaYUC9-1* and *TaYUC10* groups were active. These results indicated maximum expression of *TaYUC9-A1* or *TaYUC9-D1* and *TaYUC10-D* but suggested significant expression of the other genome copies of both genes. Taken with the RNA-seq data, it would appear that *TaYUC9-A1*, *TaYUC9-B*, *TaYUC9-D1, TaYUC10-A* and *TaYUC10-D* are all strongly up-regulated during grain fill.

### IAA production coincides with increases in grain weight and starch content from 5 to 15 DAA

To investigate the relationship between expression of *TaTAR2* and *TaYUC* genes and the auxin content of the developing grains, we measured IAA by combined liquid chromatography-tandem mass spectrometry in multiple reaction monitoring mode (LC-MS/MS MRM) with [^13^C] IAA as the internal standard in extracts of grains from 5 to 20 DAA. As shown in Fig. 2c, the IAA content increased more than 30-fold during grain development with the largest increase occurring between 5 and 15 DAA. Figures 2d and 2e illustrate the grain fresh weight (FW) and starch content respectively, at the same developmental stages. Both parameters also increased maximally between 5 and 15 DAA, coinciding with the major increase of IAA in developing wheat grains.

## Discussion

### Phylogenetic analysis of all *TAR* and *YUCCA* genes in wheat

The TAR/YUCCA pathway has been demonstrated as the main route of IAA synthesis in plants (Mashiguchi et al. 2011; Won et al. 2011). TAR and YUCCA activity has also been reported as the major source of IAA in maize and rice cereal grains (Bernardi et al. 2012; Chourey et al. 2010; Nonhebel and Griffin 2020). However, there has been no comprehensive study of these genes in wheat. Additionally, IAA is widely cited as a key signalling component during grain development. Thus, there is a need to understand its production in this important cereal crop. Finally, the frequently cited but anomalous work on the *TGW6* gene (Hu et al. 2016a; Ishimaru et al. 2013) in wheat and rice has effectively ignored this source of IAA in proposing that the activity of an IAA-glucose hydrolase can regulate the IAA content of grains.

Here, we present the first complete list and phylogenetic analysis of YUCCA genes/proteins in wheat. We have named these genes based on their homology to rice YUCCAs, rather than in comparison to Arabidopsis proteins (as was done with the previously named TaYUC10 group). YUCCA proteins are quite divergent and there are not unambiguous orthologous relationships between cereal YUCCAs and their homologues in dicots. On the other hand, most rice YUCCAs have conserved homologues in other grasses, with bootstrap values indicating high reliability of relationships (Russell French et al. 2014). It is therefore likely that the sub-functions of YUCCAs may be conserved within the cereals. The systematic naming of all genes in a family rather than *ad hoc* naming of individual genes is important to avoid future confusion. In naming genes, we have followed the convention in the Wheat Gene Catalogue of appending the genome label to each name. In line with this we have renamed the TaYUC10 group, previously described by Li et al. (2014) as *TaYUC10-B1, TaYUC10-A* and *TaYUC10-D* with the addition of *TaYUC10-B2*.

In clade I, each of the functional rice YUCCA sequences has at least one close homologue in wheat, suggesting conservation of function. Most (OsYUC1, OsYUC3, OsYUC4, OsYUC5 and OsYUC8) have a triplet of co-orthologous proteins. However, OsYUC7 has six co-orthologues in wheat suggesting a gene duplication event in a wheat progenitor. In clade II, the situation is more complex. Both OsYUC9 and OsYUC11 have close homologues in wheat, barley and Brachypodium. This is similar to maize and sorghum (*Sorghum bicolor*) (Russell French et al. 2014). However, the YUC9 group has duplicated in wheat and Brachypodium; the TaYUC9-1 group is most similar to OsYUC9 and its expression profile is also similar to the rice gene whereas the TaYUC9-2 group is more divergent. The TaYUC10 group appears to form a separate Pooid branch that has undergone gene expansion, with multiple close homologues on separate chromosomes in wheat (4A, 5B, 5D and 6B) and Brachypodium, as well as apparent tandem repeats in Brachypodium; barley has 10 proteins in this group. The phylogeny of the group is somewhat ambiguous with low bootstrap values and different programmes (Maximum Likelihood versus Neighbour Joining) variously placing the group with OsYUC11 or OsYUC12. Careful pairwise comparison of the wheat sequences with both OsYUC11 and OsYUC12 supports the maximum likelihood tree shown, placing the group closest to OsYUC11 rather than OsYUC12 and OsYUC14. This is interesting, as one of us had previously noted OsYUC12 has orthologues in maize and sorghum with similar expression profiles (Russell French et al. 2014).

Our phylogenetic analysis of TAR proteins in wheat updates the previous study by Shao et al. (2017), with the addition of three new TaTAR2 proteins, TaTAR2.6-1Ba, TaTAR2.6-1Bb and TaTAR2.6-U. These new proteins, with TaTAR2.4-7A, TaTAR2.5-1A and TaTAR2.5-1B form a separate branch, with no orthologues in rice, barley or Brachypodium. This group of enzymes may have distinct sub-function in wheat.

### Expression of *TaTAR2.3-1B, TaYUC9-1* and *TaYUC10* correlates with increasing IAA content during grain fill from 5 to 15 DAA

Results from both qRT-PCR and analysis of RNA-seq data confirmed the observation of Shao et al. (2017) that *TaTAR2.3-1B* is the most highly expressed tryptophan aminotransferase in developing wheat grains. Strong up-regulation occurred during early grain fill between 5 and 15 DAA coinciding with a similar increase in IAA content of the grains over the same interval. This suggests that TaTAR2.3 is primarily responsible for catalysing the first step in IAA synthesis of developing grains. A similar increase in expression of orthologous genes *ZmTAR1* and *OsTAR1/OsFBL* in developing grains of maize and rice also coincides with the major increase in IAA content, indicating conservation of expression of this clade (Abu-Zaitoon et al. 2012; Chourey et al. 2010). In wheat, expression of *TaTAR2.3-1B* appears to be specific to the developing grains, with no expression detectable in other parts of the plant. This may represent additional subfunctionalisation due to the larger number of TAR genes in wheat compared to other cereals.

The expression of *TaTAR2.2-1A, TaTAR2.2-1D* and *TaTAR2.5-1B* at 2 DAA, followed by their rapid down-regulation suggests IAA produced during very early grain development is regulated separately from that during the grain fill period. This is similar to rice where *OsTAR2* appears to be the dominant gene expressed at 1 DAA, whereas *OsTAR1* is expressed later (Abu-Zaitoon et al. 2012). The promotive effect of TAR and IAA for grain fill in wheat is supported by the positive effect on grain yield of *TaTAR2.1-3A* overexpression (Shao et al. 2017). In addition, a mutant of the orthologous gene, *tsg1,* has a negative effect on grain size in rice (Guo et al. 2019).

Wheat genes positioned with *OsYUC1-8* in clade I are mostly expressed in vegetative and/or floral tissues with little or no expression in grains. *OsYUC1-8* are also active in similar tissues of rice plants with low expression in grains (Abu-Zaitoon et al. 2012; Yamamoto et al. 2007; Zhao et al. 2013). On the other hand, clade II genes *TaYUC9-A1*, *TaYUC9-B*, *TaYUC9-D1*, *TaYUC10-A* and *TaYUC10-D* are all highly expressed during grain fill. These results confirm data from Li et al. (2014) relating to *TaYUC10-D* (previously *TaYUC10.3*) but expression of *TaYUC9-A1*, *TaYUC9-B* or *TaYUC9-D1* during grain fill has not been described previously. The importance of clade II YUCCA for the production of IAA in cereal grains is demonstrated by similar data from maize and rice (Abu-Zaitoon et al. 2012; Chourey et al. 2010). Interestingly, ZmYUC1 appears to be primarily responsible for IAA production in developing maize grains (Chourey et al. 2010), whereas three genes, *OsYUC9, OsYUC11* and *OsYUC12* are all up-regulated in rice grains. These rice genes have differences in their expression profiles, with earlier and more restricted up-regulation of *OsYUC12* (Nonhebel and Griffin 2020). A similar situation may occur in wheat; *TaYUC9-1* genes were maximally up-regulated between 10 and 15 DAA and remained active at 20 DAA, similar to the orthologous *OsYUC9* (Nonhebel and Griffin 2020). Expression of the *TaYUC9-1* group also increased in parallel with *TaTAR2.3-1B* and correlated most closely with the increasing IAA content of grains suggesting that these genes may be primarily responsible for IAA production during grain fill and similar again to the situation in rice. On the other hand, *TaYUC10* was up-regulated earlier than *TaYUC9-1* with a maximum increase between 5 and 10 DAA, and a decline in expression at 20 DAA. This expression profile is similar to that of *OsYUC12* in rice. Although *TaYUC10* and *OsYUC12* are not directly orthologous, their expression profile distinct from that of *TaYUC9-1* suggests a requirement for very precise and localised IAA production regulated by separate genes during early grain fill, as suggested by Nonhebel and Griffin (2020).

### Localisation of IAA production at 15 DAA in developing wheat grains

IAA production occurs mostly in endosperm of rice (Abu-Zaitoon et al. 2012; Russell French et al. 2014) and maize (Bernardi et al. 2019; Chourey et al. 2010). Forestan et al. (2010) observed high expression of the auxin transporter gene *ZmPIN1* in the periphery of maize endosperm and the embryo surrounding region. Furthermore, treatment with the auxin transport inhibitor, naphthylphthalamic acid (NPA) during early embryogenesis caused developmental abnormalities in the maize embryo. It is widely assumed, therefore, that the endosperm is the major source of IAA for the early embryo. Our results for wheat indicate that *TaTAR2.3-1B* and the *TaYUC10* group are similarly expressed only in wheat endosperm. However, we found high expression of *TaYUC9-1* in the embryo at 15 DAA, with lower expression in the endosperm. As this result was novel, the experiment was carefully checked and repeated. The observation was also confirmed by RNA-seq data from a different variety of wheat. It is probable therefore, that the embryo is only dependent on IAA imported from the endosperm at the very early stages of development.

Several reports suggest IAA may be important in aleurone layer and transfer cell development and crucial for nutrient uptake into the grains. Forestan et al. (2010) showed high immunolocalisation signal of IAA in the aleurone of maize kernels and use of NPA interfered with normal development of the aleurone. Recent work by Bernardi et al. (2019) suggests IAA has a role in the formation of the basal endosperm transfer cell layer (BETL) in maize. This is supported by the poor grain fill of auxin-deficient *de18* and *dek18* mutants of maize (Bernardi et al. 2012; Bernardi et al. 2016). Interestingly, *TaYUC-9.1* genes are most highly expressed in the transfer cells and aleurone layer of wheat grains, with lower expression in the starchy endosperm. This indicates the importance of IAA in these key cell layers is conserved between cereals and warrants further direct investigation.

### IAA production by TAR/YUCCA versus grain weight genes *TaTGW6* and *TaTGW-7A*

Our results have demonstrated high activity of the TAR/YUCCA pathway for IAA biosynthesis in developing wheat grains, coinciding with a large increase in the IAA content of grains. As in maize and rice, IAA in wheat grains accumulates particularly during early grain fill and coincides with the initiation of starch production. In other seeds, it has been proposed to initiate the differentiation of endosperm transfer cells and starch production (Bernardi et al. 2019; McAdam et al. 2017; Nonhebel and Griffin 2020). On the other hand, a negative effect of IAA on grain fill has been reported in publications relating to the *TGW6* genes in rice (Ishimaru et al. 2013) and wheat (Hu et al. 2016a), and *TaTGW-7A* in wheat (Hu et al. 2016b). Although our study did not investigate the function of these genes, it does raise some important questions. Most obviously, it is unclear how an inactive IAA glucose-hydrolase gene could have a major effect on the IAA content of grains when the TAR/YUCCA pathway is strongly up-regulated. Although no mutants of these *TaTAR2.3, TaYUC9-1* and *TaYUC10* are available in wheat to verify that they regulate IAA content, defective kernel mutants of maize with reduced ZmYUC1 activity demonstrate the importance of this pathway in cereal grain fill. Further, it is clear that there are strong similarities between IAA production in rice, maize and wheat as discussed earlier.

In addition, data on the IAA content of wheat grains with different alleles of *TaTGW6* and *TaTGW-7A* do not provide strong support for their role in IAA production either. Measurements of the IAA content of grains reported by Hu et al. (2016a) used an HPLC system with a UV absorbance detector set to 250 nm. This is an inadequate method for determining hormone contents, lacking both sufficient specificity to be sure it is measuring the correct compound as well as internal standard required to account for loss of analyte during sample work-up. In addition, the amounts of IAA found at 20 DAA, the stage at which maximum effect of the inactive genes is reported to occur, were widely variable. Values of IAA content reported in wheat varieties with the active *TaTGW6-a* allele varied from 1300– 4700 ng/g FW compared to about 800–1200 ng/g FW in varieties with the inactive or less active b and c alleles (values estimated from graphs in Hu et al. (2016a)). Similar variation was reported in the *TaTGW-7A* paper (Hu et al. 2016b). Large variation was also seen in the mean grain size from wheat varieties with active alleles of both *TaTGW6* and *TaTGW-7A*; this variation bears no relation to the IAA content. Finally, it is worth noting that neither gene in wheat has been experimentally characterised. TaTGW6 is assumed to hydrolyse IAA-glucose based on a report on the rice gene product (Ishimaru et al. 2013). Although the authors refer to *TaTGW-7A* as an “IGPS-like gene” its product is not actually a homologue of IGPS in other plants. In fact, the authors merely state that “the deduced amino acid sequences showed the presence of highly conserved TIM-br_sig_trns and UPF0261 functional domains, forming a TIM barrel fold structure, the same as IGPS”. IGPS is an essential enzyme required for tryptophan synthesis; the wheat proteome has three sequences (encoded by TraesCS2A02G335200, TraesCS2D02G329500, TraesCS2B02G348500) with between 54% and 56% amino acid identity to the experimentally characterised Arabidopsis IGPS (Li et al. 1995). These genes are expressed in the wheat plant including grains for the purpose of tryptophan production. None of these wheat sequences nor the Arabidopsis IGPS encoded by AT2G04400 has significant homology to *TaTGW-7A,* which is very unlikely to be an indole-3-glycerol phosphate synthase.

## Conclusion

In this study, we have demonstrated that auxin biosynthesis genes *TaTAR2.3-1B, TaYUC9-A1, TaYUC9-B, TaYUC9-D1, TaYUC10-A* and *TaYUC10-D* are highly active in wheat grains from 10 to 20 DAA. Their expression correlates positively with IAA levels, grain weight and starch synthesis during major grain fill period in wheat. The gene expression data combined with accurate measurements of grain IAA content from 5 to 20 DAA suggests the TAR/YUCCA pathway is a major source of auxin in developing wheat grains. This raises serious questions about how a mutation in *TGW6*, encoding a putative IAA-glucose hydrolase could have the reported effects on grain IAA content. Our data on localisation of auxin biosynthesis gene expression confirm that IAA is produced in wheat endosperm as in other cereals. In addition, the novel finding of *TaYUC9-1* activity in embryos at 15 DAA suggests that embryos have their own source of IAA by this stage of development. Differences in the expression profiles of *TaYUC9-1* and *TaYUC10* groups suggest some sub-functionalisation. The expression of *TaYUC9-1* genes in aleurone and transfer cells corroborates the suggestion by Bernardi et al. (2019) that IAA may be involved in differentiation of these cells in maize. This may partly explain the importance of IAA for grain fill.

## Declarations

### Availability of data and material

Not applicable

### Code availability

Not applicable

### Author contribution statement

MRK and HMN conceived and designed the research. MRK performed all the experiments. DB contributed to growing wheat plants and preparation of manuscript. GW helped in analysing gene expression data. MRK and HMN wrote the manuscript. All authors read and approved the manuscript.

### Key message

Expression of *TaTAR2.3*, *TaYUC9-1* and *TaYUC10* coincides with increasing IAA content, grain weight and starch content from 10-15 DAA, highlighting the importance of the TAR/YUCCA pathway in wheat grain filling.

## Acknowledgements

The authors are grateful to Kirsten Drew for the analysis of IAA. The authors also acknowledge the Australian Government for providing a Research Training Program (RTP) PhD scholarship to Muhammed Rezwan Kabir.

