## Supplementary material for "Expression of *TaTAR2.3-1B*, *TaYUC9-1* and *TaYUC10* correlates with auxin and starch content of developing wheat grains"

**Supplementary Table S1.** Primers used for expression analysis of *TaTAR2* and *TaYUC* genes

| Gene group | Primer sequence | Product size (bp) |
|---|---|---|
| *TaTAR2.1-3A*<br>*TaTAR2.1-3B*<br>*TaTAR2.1-3D* | F: CTACTACTCGTCGTACCCTGCC<br>R: CGGTGAAGAGCATGATGTCG | 251 |
| *TaTAR2.2-1A*<br>*TaTAR2.2-1B*<br>*TaTAR2.2-1D* | F: CGACCATGTTTGAGGCCTTC<br>R: CCATGAACCAGCACACGTTA | 114 |
| *TaTAR2.3-1A*<br>*TaTAR2.3-1B*<br>*TaTAR2.3-1D* | F: TCATGCTCTTCACCGTGTCA<br>R: GCTTTGCTCCATGAACTTGC | 113 |
| *TaTAR2.4-7A* | F: CACATGCATCGTCTCATCCC<br>R: ACCACTGCCTTATTGCTCCT | 122 |
| *TaTAR2.5-1A*<br>*TaTAR2.5-1B* | F: ATCAACCTCGAGCTCGGC<br>R: CTGGGCATTGGAGAAGTAGC | 117 |
| *TaTAR2.6-1Ba*<br>*TaTAR2.6-1Bb*<br>*TaTAR2.6-U* | F: TTAGAAGAAGGGTGACGCGT<br>R: GTCGCCGGTTTATGTGGTAC | 186 |
| *TaYUC9-A1*<br>*TaYUC9-B*<br>*TaYUC9-D1* | F: ACCAAGGAGATCTGGAACGT<br>R: GGGGTGTGGATCTTCATGGA | 164 |
| *TaYUC9-A2*<br>*TaYUC9-D2* | F: ACGAGGGAGATCATGAACGC<br>R: GATCTTGTCGTAGGTGCCGA | 204 |
| *TaYUC10-A*<br>*TaYUC10-B1*<br>*TaYUC10-B2*<br>*TaYUC10-D* | F: GTCCGATTCATGTAATGACAAAG<br>R: GGTGCCAACATCAATCACTG | 209 |
| *TaYUC11-A1*<br>*TaYUC11-A2*<br>*TaYUC11-D* | F: CCAGAGATCGTTGGCCTACA<br>R: CCCACATCCAACAACAAGCA | 108 |

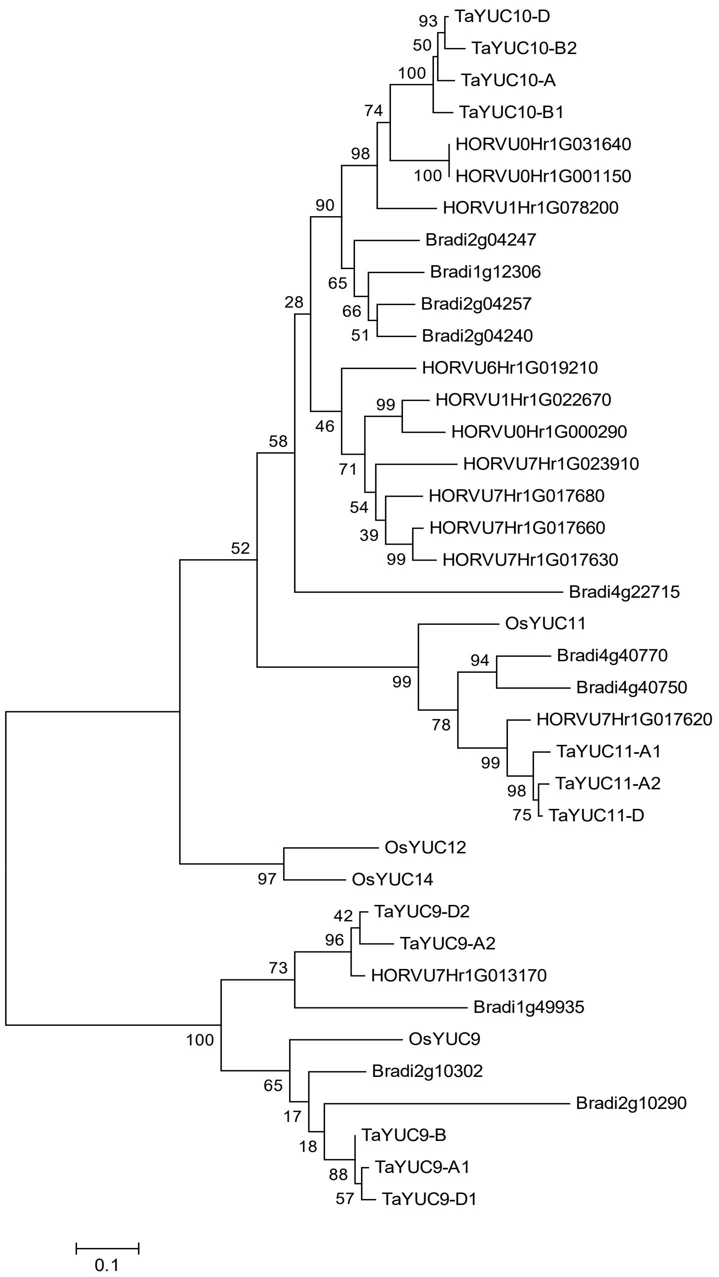


**Supplementary Figure S1.** Phylogenetic tree showing relationships between Clade II YUCCA proteins from *Triticum aestivum* (TaTAR), *Oryza sativa* (OsTAR), *Brachypodium distachyon* (Brad) and *Hordeum vulgare* (HORVU). The tree was constructed in MEGA7.0.26 (Kumar *et al.*, 2016) using Maximum Likelihood method (Jones *et al.*, 1992). Multiple sequence alignment was performed by MUSCLE (Edgar, 2004). Bootstrap confidence levels were obtained using 500 replicates (Felsenstein, 1985). Evolutionary distances were computed using Poisson correction method (Zuckerkandl and Pauling, 1965). Scale bar=0.10, amino acid substitutions per site
